# Highly Extensible Physically Crosslinked Hydrogels for High-Speed 3D Bioprinting

**DOI:** 10.1101/2024.08.05.606733

**Authors:** Ye Eun Song, Noah Eckman, Samya Sen, Carolyn K. Jons, Olivia M. Saouaf, Eric A. Appel

## Abstract

Hydrogels have emerged as promising materials for bioprinting and many other biomedical applications due to their high degree of biocompatibility and ability to support and/or modulate cell viability and function. Yet, many hydrogel bioinks have suffered from low efficiency due to limitations on accessible printing speeds, often limiting cell viability and/or the constructs which can be generated. In this study, we report a highly extensible bioink system created by modulating the rheology of physically crosslinked hydrogels comprising hydrophobically-modified cellulosics and either surfactants or cyclodextrins. We demonstrate that these hydrogels are highly shear-thinning with broadly tunable viscoelasticity and stress-relaxation through simple modulation of the composition. Rheological experiments demonstrate that increasing concentration of rheology-modifying additives yields hydrogel materials exhibiting extensional strain-to-break values up to 2000%, which is amongst the most extensible examples of physically crosslinked hydrogels of this type. We demonstrate the potential of these hydrogels for use as bioinks by evaluating the relationship between extensibility and printability, demonstrating that greater hydrogel extensibility enables faster print speeds and smaller print features. Our findings suggest that optimizing hydrogel extensibility can enhance high-speed 3D bioprinting capabilities, reporting over 5000-fold enhancement in speed index compared to existing works reported for hydrogel-based bioinks in extrusion-based printing.

## 1. Introduction

Three dimensional (3D) bioprinting has revolutionized tissue engineering technology by enabling the fabrication of tissue structures in controlled shapes and placement of living cells or cell aggregates in precise layouts.^[1]^ Bioprinting allows automated construction of functional tissue scaffolds through layer-by-layer deposition of bioinks, generally made of biomaterials, leading to high-throughput production of artificial tissue scaffolds. Bioprinting technologies can be categorized into three types depending on the method of deposition and patterning: inkjet, micro-extrusion, and laser-assisted printing.^[2]^ Specifically, micro-extrusion methods are extensively utilized due to their high versatility and compatibility with diverse gelation methods as well as general accessibility and affordability. Yet, the main challenges in micro-extrusion bioprinters are typically low cell viability, low print resolution, and limited printing speed.

Advancement of bioprinting technology in the recent decade was aligned with the improvement of biomaterials and bioinks.^[3]^ Hydrogels - water swollen 3D polymer networks – represent a highly studied class of bioinks with excellent biocompatibility and tunable chemistry enabling facile engineering for cell attachment, growth, migration, and differentiation, as well as effective protection from mechanical disruption due to high shear stresses during injection or processing.^[4-8]^ Recent developments in hydrogel bioinks has addressed many crucial challenges in bioprinting. For example, jammed microgel bioinks comprising cross-linked hydrogel microparticles that are aggregated together by confinement can be formed from materials typically incompatible with bioprinting in their bulk form and exhibit excellent extrudability while improving the viability of entrapped cells on account of the increased porosity within the systems.^[9-10]^ Moreover, double-network hydrogel bioinks from dual crosslinking exhibit tunable mechanical strength while maintaining good cell viability, cell proliferation, and structural integrity.^[11-13]^ Beyond synthetic hydrogel systems, protein-based hydrogels have also been widely studied and are promising candidates for bioinks providing cell encapsulation environments which highly resemble living tissues.^[14-16]^ Various multicomponent hydrogel-based bioinks, which comprise more than one type of polymer or hydrogel network to achieve combined benefits of single component bioinks, have shown advanced mechanical stability for shape fidelity post printing while maintaining good biofunctionality.^[17]^ Yet, while many bioinks have become significantly more advanced with regard to cell viability and mechanical properties, limitations in printing speed have received much less attention and have not yet been adequately addressed.

During printing of filaments at higher speeds, bioinks experience large stretching strains and forces. When this stretching strain surpasses the extensional strain of the bioink material, the printed features of the filament display discontinuities and irregular print diameters.^[18, 19]^ Hence, high extensibility of the filament plays a key role in improving the printing resolution at increased speed.^[20]^ Hydrogel-based bioinks generally suffer to obtain high structural integrity and shape fidelity due to their high water content, making it challenging to achieve high printability in general.^[53]^ Moreover, very few hydrogel materials exhibit sufficient extensibility to achieve high printability at high-speed printing, and indeed, very few studies have comprehensively evaluated the impact of bioink extensibility in the context of bioprinting.^[20, 21]^

In this work, we report the development of a broadly tunable hydrogel platform with extreme extensibility (upwards of 2000%), prepared by simple mixing of hydrophobically modified cellulosic biopolymers and common biopharmaceutical drug product excipients including surfactants and cyclodextrins. We demonstrate that alteration of the hydrogel composition enables control over the crosslink dynamics for these physically crosslinked hydrogels, imparting control over shear-thinning and dynamic mechanical properties such as viscoelasticity, stress relaxation, and extensibility. We analyze the cell compatibility of formulated hydrogels and characterize the relationship between bioink extensibility and printability at high speeds (5300-fold improvement in achievable velocity factor *S* = printing velocity over material flow velocity) using an extrusion-based 3D printer.

## 2. Results and Discussions

### 2.1. Formulation of hydrogel

It has been widely reported that modification of polysaccharides with long-chain hydrophobic moieties can enable robust, non-covalent, hydrophobic interactions between polymer chains in aqueous media that yields hydrogel-like materials.^[22, 23]^ Indeed, numerous commercial products based on this class of materials exist, and they are used in a wide variety of applications from food products to cosmetics to hydraulic fracturing, where they are typically used as viscosity modifiers.^[24-26]^ With some materials, such as hydroxypropyl methylcellulose stearoxy ether (HPMC-C18; available commercially as Sangelose^®^ from Daido chemical corporation) dissolving in aqueous media immediately forms robust, colorless hydrogels (**Figure 1a**). In aqueous media, the micellization of the stearyl groups pendant from the HPMC backbone upon dissolution facilitates the facile formation of a robust physical hydrogel network. While the micelles serve as reversible, physical crosslinks between the HPMC polymer chains, these crosslinking interactions exhibit very slow exchange dynamics on account of the long, saturated, and highly hydrophobic stearic chains. It was previously reported that addition of cyclodextrins can change the viscosity of HPMC-C18 polymer solutions and gels.^[27]^ It was speculated that the cyclodextrin binds the stearyl side chains on the HPMC-C_18_ polymers, thereby weakening the associations between the hydrophobic stearyl chains responsible for the network structure. We hypothesized that other chemical additives which interact with the stearyl groups on the HPMC-C_18_ polymers could tune the exchange dynamics of the micellar crosslinks of these materials, for instance by introducing free volume into the micellar structure, enabling refined modulation of the hydrogel properties that could enable development of bioinks with useful properties for high-speed printing applications. In this work, we assessed the impact of different additives on the dynamic mechanical properties and flow properties of HPMC-C_18_ based hydrogel materials (**Figure 1b**), verifying our hypothesis through detailed rheological experiments, and evaluated the use of these materials as bioinks.

**Figure 1.**
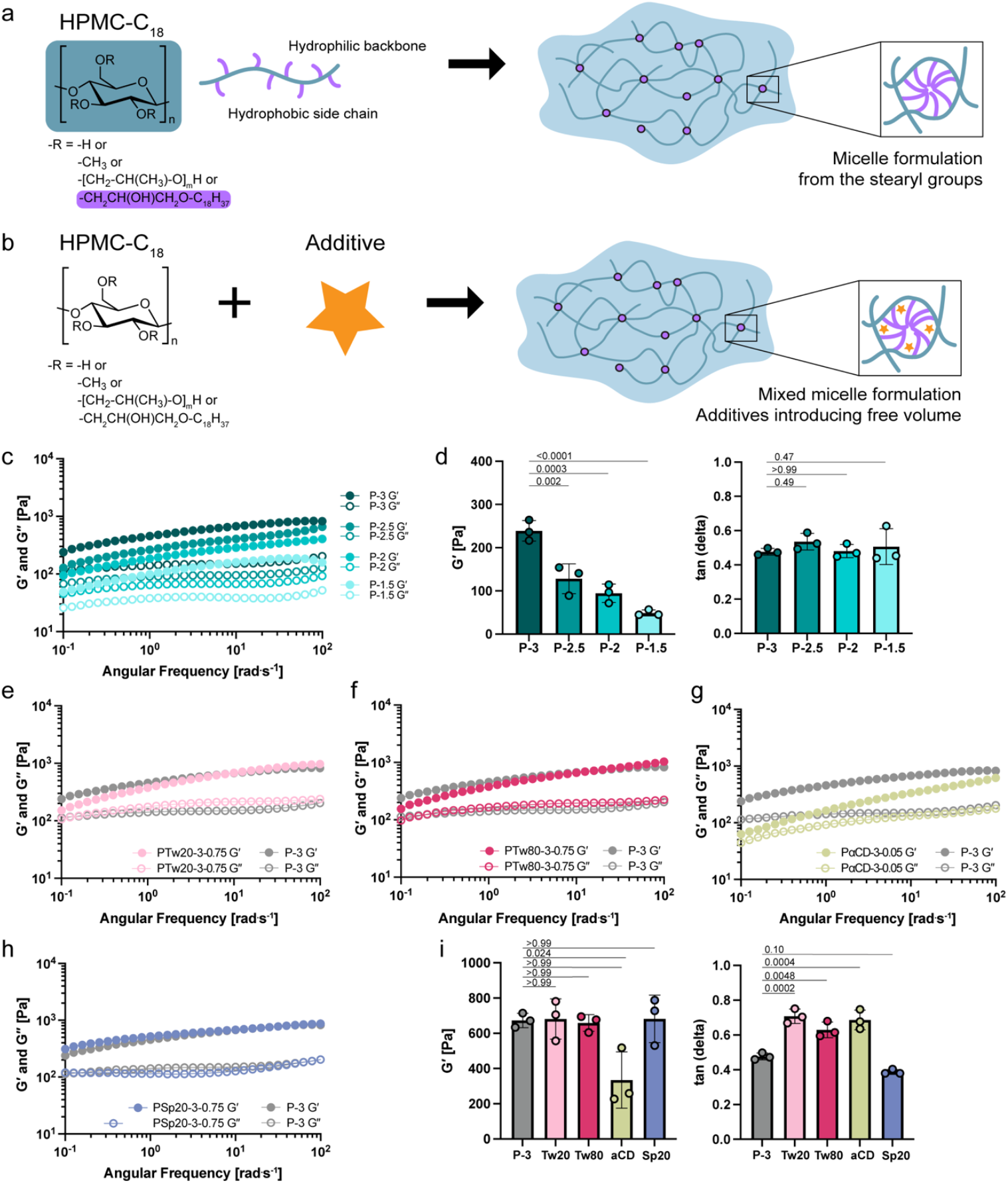
(a) Schematic illustration of HPMC-C_18_ hydrogel formulation upon mixing HPMC-C_18_ in a solvent. (b) with and without additives. Additives tune the dynamic of micelle crosslinks by introducing the free space. (c) Frequency sweep of HPMC-C_18_ hydrogels formulated at different weight percent. (3wt%, 2.5wt%, 2wt%, and 1.5wt%). (d) Storage modulus (*G*′) and tan(*δ*) values read at 10^−1^ rad.s^-1^ from (c). Changing the polymer weight percent can tune the stiffness but not the viscoelasticity of the gel. (e – h) Frequency sweep of HPMC-C_18_ hydrogel with different additives: Tween 20, Tween 80, alpha-cyclodextrin, and Span 20. Weight percent of each component is noted after the component name. (e.g., PTw20-3-0.75 refers to the hydrogel formulation with 3 wt % of HPMC-C_18_ polymer and 0.75 wt % of Tween 20.) (i) *G*′ and tan (*δ*) (defined as *G*′′/*G*′) values read at 10 and 10^−1^ rad.s^-1^ each from (e – h). HPMC-C_18_ hydrogels with additives can tune the viscoelasticity of the gel. One-way ANOVA was used for multiple comparisons.

### 2.2 Shear rheological characterization

To quantify the linear viscoelasticity of the hydrogel formulations, we made four hydrogel samples titrating HPMC-C_18_ polymer content and performed various shear rheometry tests (**Figure 1c-i**). Hydrogel samples are denoted P (e.g., P-3), followed by a number that indicates the weight percent of the HPMC-C_18_ polymer in formulation. Any additives in the hydrogel will be indicated after P, and the weight percent of the additive will be marked after the weight percent of the polymer. For example, a hydrogel sample named PTw20-3-0.75 contains 3 wt% of the HPMC-C_18_ polymer and 0.75 wt% of Tween 20 as an additive. Hydrogels made with HPMC-C_18_ polymer will be referred to as HPMC-C_18_ hydrogels. Frequency sweeps were performed to examine the viscoelastic response over multiple timescales. All formulations display solid-like rheological behavior, showing higher storage (*G*′) over loss (*G*′′) moduli across the entire range of observed frequencies (0.1 to 100 rad.s^-1^). Increasing the polymer content increases the stiffness of the hydrogels (**Figure 1d**). However, titrating the polymer content has minimal effect on the slopes of the storage and loss moduli curves. For all formulations, a crossover between the shear storage and loss moduli, the frequency at which represents the inverse of the hydrogel network relaxation time, is not observed in the measured frequency range and is expected to be visible at longer time (low frequencies) since the slopes of both shear moduli are similar within the measured frequency region. Increasing the polymer content also largely preserves the viscoelasticity (relative solid-fluid nature of the network, represented by tan (δ)) of the hydrogel and primarily tunes the stiffness (*G*′) (**Figure 1d**). An increase in apparent yield stress is also observed as the polymer content is increased (**Figure S1**).

To explore the effect of additives on the dynamic mechanical properties of the HPMC-C_18_ hydrogels, we formulated hydrogel samples with additives by simply mixing HPMC-C_18_ polymer and a choice of additive. We screened non-ionic surfactants with high and low hydrophilic-lipophilic balance (HLB) values and alpha cyclodextrin (αCD) as additives and observed changes in the mechanical properties of hydrogels. We chose to screen polysorbates (Tweens) and sorbitan esters (Spans) for non-ionic surfactants with high and low HLB values since both surfactants are FDA approved drug products that are commonly used in biologic drug product formulations, cosmetics, food, and agricultural applications. In all formulations, the amount of the HPMC-C_18_ polymer and the additives remained constant unless mentioned otherwise.

Tween surfactants are non-ionic surfactants with high HLB values (15.0 – 16.7), hence favoring solubility in aqueous buffers.^[28, 29]^ Tween surfactants were studied to inspect the effect of hydrophilic surfactants on the mechanical properties of HPMC-C_18_ hydrogels (**Figure 1e, 1f, and S2**). We first compared how the change in the tail length of the surfactant affects the rheological properties of the hydrogel by comparing Tween 20 (Tw20), Tween 40 (Tw40) and Tween 60 (Tw60) with laurate (C12), palmitate (C16), and stearate (C_18_) tails respectively. Hydrogel formulation with Tw20 exhibits a change in the slope of *G*′, and a crossover between the storage and loss moduli shifts to the right (higher frequencies) accordingly (**Figure 1e**). Notice that the addition of Tw20 increased tan(*δ*), i.e. increased the fluid-like characteristics relative to solid-like, but overall maintained a solid, or gel-like response since tan(*δ*) < 1 even with surfactants, but largely did not affect *G*′ values approaching the high-frequency plateau, which remained around 700 Pa (**Figure 1i**). Since the crossover between the storage and loss moduli represents a relaxation time of the hydrogel network, a rightward shift in the crossover point means a shorter relaxation time is observed in the hydrogel when Tw20 is added (i.e., more dynamic network). We also confirmed this numerically by fitting the frequency sweep data to a continuous relaxation spectrum (**Figure S3**). The change in gel viscoelasticity (tan(*δ*)) and the crossover point (relaxation time) is more prominent in shorter tail surfactants, as shown in increasing tan(*δ*) values and progressively small changes in the slope of *G*′ of the hydrogel formulations with Tw40 and Tw60. Specifically, the stearic acid tail of Tw60 had no significant effect in changing the rheological properties of the HPMC-C_18_ hydrogel (**Figure S2**). Similar changes in the relaxation dynamics and gel viscoelasticity from Tw20 sample were examined in the formulation with Tw80 (**Figure 1f and 1i**). The unsaturated fatty acid tail (i.e., oleate) of Tw80 surfactant created more dynamic crosslinks than fully saturated fatty acid tail (i.e., stearate) of Tw60. The effect of surfactants with multiple tails was also investigated using Tween 65 (i.e., three stearate tails) and Tween 85 (i.e., three oleate tails). Neither surfactant had any significant effect on gel dynamics and viscoelasticity (**Figure S2**).

In summary, adding Tween surfactants tuned the gel viscoelasticity while not significantly changing the stiffness of the hydrogel. We hypothesize that adding Tween surfactants alters the surface properties of the noncovalent crosslinking micelles, enabling tuning of gel dynamics and viscoelasticity without changing the stiffness of the hydrogel. Upon the addition of αCD to HPMC-C_18_ hydrogels, we observed similar shift in the frequency crossover to the right (i.e., more dynamic) while the modulus value approaching the apparent high-frequency plateau remained relatively unchanged (**Figure 1g**). The crossover shift, however, was greater with αCD than with Tween surfactants even at a lower weight percent of αCD. We hypothesize that the inclusion of αCD may disrupt the formation of the micelle crosslinks, since the αCD cavity can bind strongly to the stearoyl group of HPMC-C_18_.^[25]^

Span surfactants are non-ionic surfactants with low HLB values (4.3 – 8.6) and favor solubility in lipophilic or hydrophobic solvents.^[29]^ Span 20 (Sp20), Span 40 (Sp40), and Span 60 (Sp60), with C12, C16, and C_18_ tails respectively, were examined to study the change in hydrogel rheological properties due to hydrophobic surfactants additives with different tail lengths. Since we initially hypothesized that the additives that interact with the stearic group of HPMC-C_18_ will tune the crosslink dynamics, Spans were not anticipated to sufficiently modulate hydrogel properties due to their hydrophobicity. As expected, Sp20 had almost no effect in tuning the viscoelasticity of the gel (**Figure 1h**). Sp40 and Sp60 also did not change the rheological properties of the HPMC-C_18_ hydrogel significantly (**Figure S4**).

After screening the above series of surfactant additives, we characterized the linear viscoelasticity and nonlinear yielding properties of the HPMC-C_18_ hydrogels at increasing concentration of additives of interest. We chose to proceed with Tw80 given its common use in biomedical and pharmaceutical areas.^[30, 31]^ A frequency sweep reveals that the crossover between *G*′ and *G*′′ shifts gradually to the right (higher frequencies) as the concentration of Tw80 increases in HPMC-C_18_ hydrogel formulations (**Figure 2a**). This tendency is also confirmed by the decreasing relaxation time with increasing Tw80 content (**Figure 2d**). We observed that the stress relaxation time decreased by 2.6 fold with the addition of only 0.1wt % of Tw80. We confirmed that the hydrogel stiffness and gel viscoelasticity can be tuned independently by controlling the concentration of each component in HPMC-C_18_ hydrogel (**Figure 2c**). While tan(*δ*) increased with Tw80 concentration, *G*′ values remained largely constant around the high-frequency plateau, which itself can be tuned by the concentration of HPMC-C_18_ polymer content as mentioned previously. Yielding behaviors of HPMC-C_18_ hydrogel formulations were analyzed with amplitude sweeps in large amplitude oscillatory shear tests at a constant frequency of 0.1 rad.s^-1^, where the crossover between the storage and loss moduli gives an estimated location of the yielding stress or strain (**Figure 2b and S5**). HPMC-C_18_ hydrogels with no additives all exhibited yielding at high strains (>300%). Addition of Tw80 reduced the strain-to-yield, mainly due to decreased *G*′ values while *G*′′ values showed negligible change. We also conducted steady-state shear flow tests varying shear rates from low to high to measure the static yield stress (**Figure 2e**). The yield stress of the hydrogel decreased significantly at 1wt% or higher Tw80. In short, these findings propose a simple method to precisely tune the stiffness, viscoelasticity, apparent yield stress, and the relaxation dynamics of the hydrogel independently by controlling the polymer and additive compositions.

**Figure 2.**
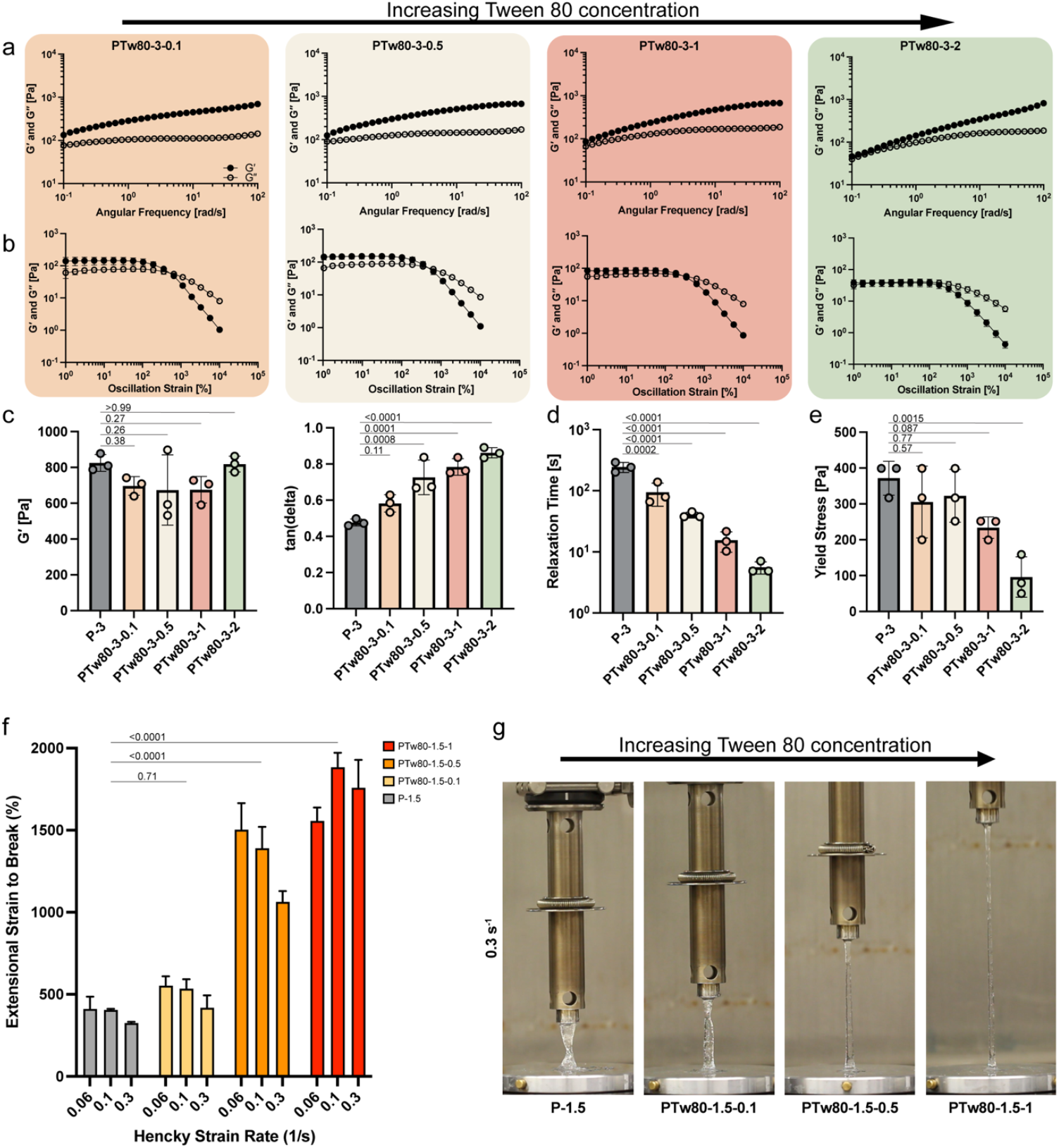
(a) Frequency sweeps of PTw80 hydrogels at different weight percent of Tw80. The crossover between *G*′ and *G*′′ shifts to the right as the hydrogel sample contains higher weight percent of Tw80. (b) Amplitude sweeps of PTw80 hydrogels with different weight percent of Tw80 measured at a frequency of 10^−1^ rad.s^-1^. (c) *G*′ and tan(*δ*) values measured at 10^2^ rad.s^-1^ and 10^−1^ rad.s^-1^ from (a), respectively. (d) Relaxation time is calculated by fitting the frequency sweep with the continuous relaxation spectrum. Gel relaxation time decreases as the weight percent of the T80 increases. (e) Yield stress is determined through stress-controlled flow sweep with increasing stress. (f) Extensional strain to break measurement of hydrogel samples with 1.5 wt% of HPMC-C_18_ polymer with different T80 contents measured at three different Hencky strain rates. (n=3) (g) Representative images of HPMC-C_18_ hydrogels with Tw80 at an extensional strain rate of 0.3 s^-1^. One-way ANOVA was used for multiple comparisons.

### 2.3 Extensional strain-to-break measurement

Displaying high extensibility is important characteristic for bioinks since they undergo high stretching strain during high-speed printing. Hence, we examined the extensibility of HPMC-C_18_ hydrogels. In previous study, yield stress fluids made from cellulose-based polymer have exhibited remarkably increased extensibility as the relaxation time decreased.^[21]^ Given the decreased relaxation time with increased additive concentration in HPMC-C_18_ hydrogels, we hypothesized that hydrogels with higher Tw80 concentration will express higher extensibility. To verify our hypothesis, we performed filament stretching extensional rheometry (FiSER) tests, a method in which a sample is located between two coaxial parallel plates while a nominally assumed uniaxial extension flow is applied to the samples as they get stretched between the plates.^[32, 33]^ We measured the extensional strain-to-break metric, which is defined as the percentage of the applied nominal strain (engineering strain) at filament breakup under an applied, constant, nominally true (Hencky) strain rate during FiSER, and this is used as the measure of extensibility in the hydrogels. We screened a variety of Hencky strain rates to detect any rate-dependent extensional behavior.^[34]^ To align with previous studies, the aspect ratio of the cylindrical hydrogel samples, given by the ratio of the initial distance between two plates to the radius of the plate, was kept at unity for all experiments.^[33]^

HPMC-C_18_ hydrogels achieved impressively increased extensibility in formulations with higher concentration of Tw80 (**Figure 2f**). Hydrogels without Tw80 exhibited strain-to-break values of approximately 400%. While the change in strain-to-break was minimal at the lowest concentration of Tw80 tested (PTw80-1.5-0.1), values increased to 2000% (20 times the initial strain) in PTw80-1.5-1 formulations, exceptionally high compared to other physically crosslinked hydrogels previously reported in the literature.^[21, 33]^ All of the hydrogel samples displayed strain rate dependent behavior. While PTw80-1.5-1 showed the lowest strain-to-break extension at low Hencky strain rate with increasing extensibility with escalating strain rate, three other formulations showed inversely related behavior between the extensional strain-to-break and strain rate. We also conducted static release assays of Tw80 to ensure that Tw80 is not released from the hydrogel network after it is formed for two hydrogel formulations (PTw80-1.5-0.5 and PTw80-1.5-1) with significantly improved extensional strain-to-break (**Figure S6**). Over 90% of Tw80 was retained in the hydrogel network for the first 24 hours. We also confirmed that less than 5% of the total Tw80 content was released additionally when left for 7 days.

Additionally, we observed that these Tw80 mixed HPMC-C_18_ hydrogels demonstrated uniformly stretching filaments (approximately uniform diameter axially, canonically homogeneous uniaxial extension) during extension rather than local thinning at a single location (**Figure 2g and Video S1**). In general, during the extension of a material, local necking and thinning is followed by further stretching of the thinner part since the stress is highest at the axial location with the lowest surface area for a given tensile force that is constant throughout the filament. This gives rise to runaway thinning and eventual failure of the filament. Localized thinning is commonly observed in dilute polymer melts or yield stress fluids.^[21, 33, 35]^ Our hydrogels showed approximately uniform (homogeneous) extension, suggesting a self-healing behavior during extension which counteracts further thinning and stress localization and resists possible deformation.^[36]^ Uniform extension is directly linked to the uniformity of filament cross section, which is a desirable feature of the 3D printed filaments.^[37, 38]^ Hence, we have created a highly extensible, uniformly stretching hydrogel system with facile formulation *via* simple mixing of polymer and a surfactant. Having confirmed the following desirable features: ease of fabrication, high extensibility, and uniformity of filaments under stretching forces, we decided to further test the potential of HPMC-C_18_ hydrogels as bioinks.

### 2.5 3D printing with PTw80 hydrogels

Prior to assessing the printability, we conducted a study of cell viability in the HPMC-C_18_ hydrogels to ensure their suitability as bioinks.^[1, 3]^ Preliminary evaluation of cell viability was done in plain HPMC-C_18_ hydrogel (P-1.5) without Tw80 through a live/dead assay after overnight culturing (**Figure S7**). In the plate control, flattened shapes of the cells were apparent which are often observed in cell work in two-dimensional stiff materials such as polystyrene.^[39]^ Cells in P-1.5 were absent with flattening and elongation of cell shapes, typical of cells cultured in two-dimensional hydrogel film with low stiffness (below 1kPa) or three-dimensional hydrogel platform.^[40, 41]^ We performed the same live/dead assay with PTw80 hydrogels since Tween surfactants have been reported of cytotoxicity in past *in vitro* studies.^[42, 43]^ We tested three formulations with Tw80, all of which displayed good cell viability. We also analyzed the cell viability post extrusion through needle to mimic the environment that cells experience during printing (**Figure S8**). While the cell viability assays on our hydrogel platform demonstrate adequate cytocompatibility for most bioink applications, the data suggest that cell viability is potentially decreasing with increasing concentration of Tw80. Therefore, further experiments to optimize the formulation compositions will be necessary for future work aimed at leveraging other cell types in specific target applications. To evaluate the ability of PTw80 hydrogels to promote cell proliferation, we performed cell proliferation assays *in vitro* using CCK-8 assays (**Figure S9**). No significant difference was observed among the different groups during the first 4 hours of culturing. An upward trend is shown when the groups were cultured for three days with the highest OD values observed in the hydrogels with highest Tw80 concentration, suggesting that PTw80 hydrogels promote cell proliferation.

To characterize the printability of HPMC-C_18_ hydrogels, we used a lab-made extrusion-based 3D printer with mechanically controlled printing stage and dispensing system (**Figure S10**).^[44]^ We were aiming to analyze the printability of hydrogels with various extensibilities at high printing speeds to validate our hypothesis that high extensibility is crucial for improved high-speed printability. To prove this experimentally, we first defined what high-speed printing means. Filament experiences high stretching strain when the speed of the nozzle relative to the printing stage (*ν*_print_) is greater than the extrusion speed (*ν*_flow_) of the material deposition.^[18, 19,45]^ Thinning of filament diameter is observed until it eventually discontinues as the stretching strain experienced is higher than the extensional strain-to-break of the bioink. Hence, the stretching and thinning of the filament we observe at high-speed printing is defined as higher nozzle movement speed (*ν*_print_) relative to material extrusion speed (*ν*_flow_). Here, we define a dimensionless value *S* as

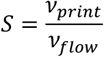

so that increased *S* would indicate an experimental condition when the filament experiences higher stretching strain due to high-speed printing. We tested four hydrogel formulations from Fig 2g at increasing *S* to analyze the printability of the filament at increasing extensibilities. Since the 3D printer we used was limited to low *ν* _print_ due to motor restrictions. We circumvented limitations in stage movement speed by controlling *ν* _flow_ (**Figure 3a**). We increased *S* by controlling *ν* _flow_ at a fixed *ν* _print_ to facilitate equivalent condition (constant extensional flow rate) to printing at varied and increased speeds (**Figure 3b**).

**Figure 3.**
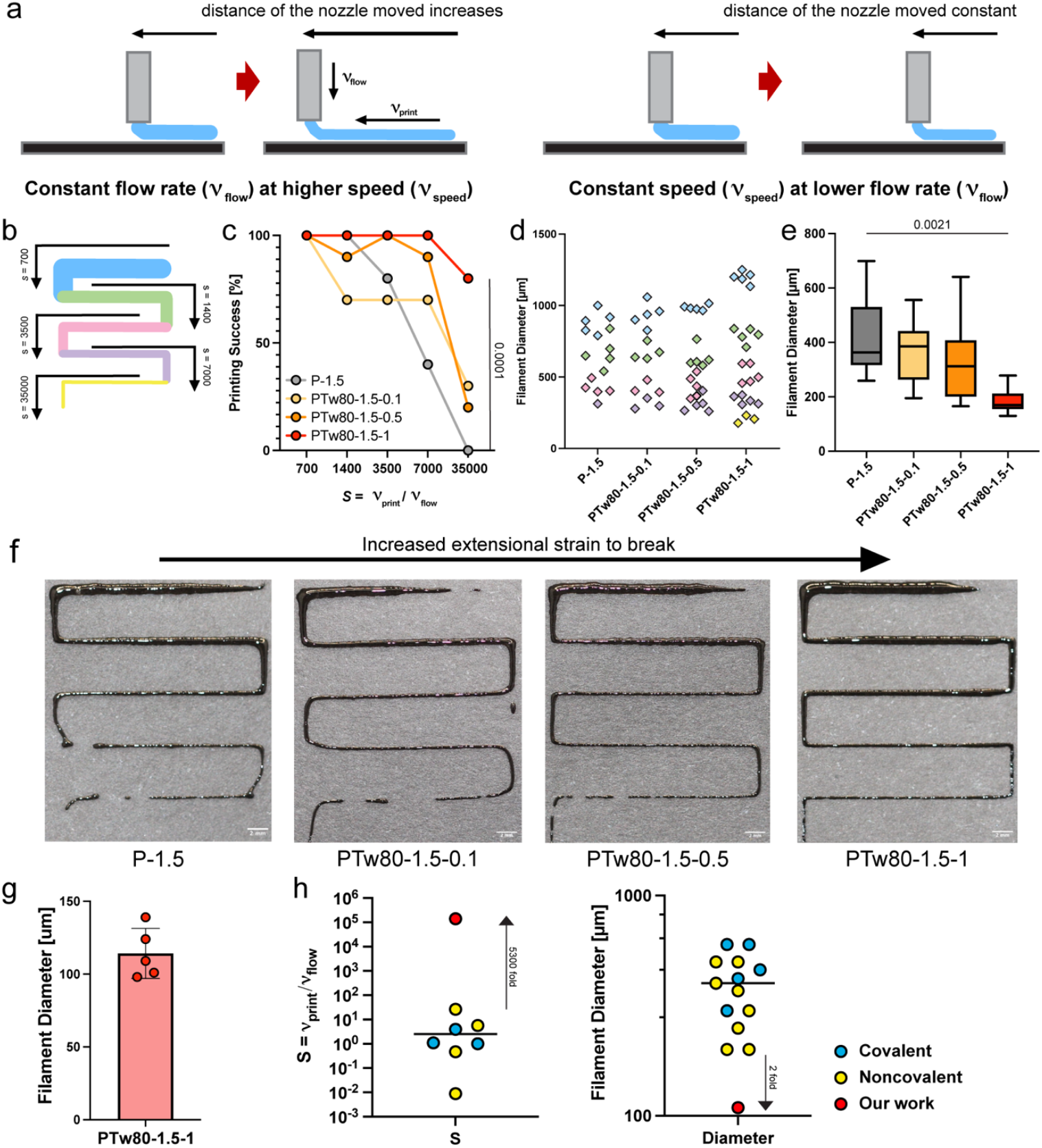
(a) Schematic illustration of filament printed while changing the speed of the nozzle and the flow rate independently. The printed filament experiences extension at a higher speed and a lower flow rate. (b) Schematic illustration of the printed filament at various value of *S*. (c) Printing success percentage plot of four different formulation of PTw80 hydrogels at various value of *S*. PTw80 hydrogels with the highest weight percent of Tw80 – highest extensional strain to break – demonstrates the best printing performance. (n = 10) (d) Average filament diameter per different value of *S* (n = 5) Color (matching with the color code in (b)) indicates the value of *S* at which the filaments were printed. (e) Minimum filament diameter measured before the break. (f) Representative images of the printed hydrogels. (g) Minimum diameter measured before the break for PTw80-1.5-1 when printed at *S* = 70000 and *S* = 140000. (Scale bar: 2mm) (h) Comparison of *S* and printed filament diameter with other hydrogel based bioinks in extrusion based 3d bioprinting (needle gauge 27G or above). Two-way ANOVA was used for printing success percentage plot in (c). One-way ANOVA was used for filament diameter plot in (e).

Filaments undergoing extrusion experience high stretching forces when the printing speed increases, or the flow rate decreases (*S* increases), and this can lead to large extensional strains and arbitrary thinning of the filament, loss of cross-sectional uniformity, and ultimately filament disconnection and failure.^[18, 19]^ Since high extensibility is hypothesized as a potential solution to mitigate filament breaking, we expected the more extensible PTw80 hydrogel to display better printability. Qualitative inspection of HPMC-C_18_ hydrogel with no Tw80 revealed typical yield stress fluid filament behavior with increasing S (decreasing flow rate) (**Figure 3f**). As the value of *S* increased by a factor of 20, filament disintegration was observed followed by the accumulation before and after the breaking. Corner shapes were not maintained, and the fidelity of individual filament shapes was compromised. Similar shapes were observed in PTw80-1.5-0.1 in which the corners were still less defined in shape and filament breaking was present at the lowest flow rate tested. Promisingly, for both PTw80-1.5-0.5 and PTw80-1.5-1 in which the extensibility increased greatly, improved printability was observed with better maintained corner shapes and filament uniformity. Our most extensible hydrogel sample, PTw80-1.5-0.1, displayed the best printed results with sharp corners and continuous, uniform filaments.

To quantify the printability of PTw80 hydrogel samples, we plotted the printing success percentage at different value of S, defining a “failure” in print as the breakage of the filaments (**Figure 3c**). HPMC-C_18_ hydrogels with no Tw80 showed a sharp decrease in printing success percentage which eventually dropped to zero. PTw80-1.5-0.1 and PTw80-1.5-0.5 showed relatively high success rate over 70%, but the percentage reduced below 30% at the highest S value. Our most extensible hydrogel candidate, PTw80-1.5-1, expressed fully successful printed filaments at all flow rates except *S* = 35000 at which the filament was still successfully printed over 80% of the total tested time. Thus, we confirmed that high extensibility is a crucial feature of bioinks to enable high speed printing without print discontinuation. To quantify the resolution of the filaments, we also analyzed the diameter of filaments printed. The average diameter of the filament was measured at different value of *S* and plotted to characterize the range of diameters that can be printed using each hydrogel samples (**Figure 3d**). The diameter of the filament will decrease as the material is subjected to greater extensibility at higher value of *S*, so we expected the filament diameters to be in separate groups of datapoints due to changing flow rates. For P-1.5 and PTw80-1.5-0.1, the datapoints are centered and less differentiated, meaning that filament thickness post printing was not varied distinctively by changing the value of *S* (either the print speed or the flow rates) due to lack of filament diameter uniformity. For PTw80-1.5-0.5 and PTw80-1.5-1 formulations, thicker filaments are plotted in more defined clusters with tighter datasets. These reveal that we can print a wider range of diameters by changing the flow rate or the print speed and precisely target the desired filament diameter with more uniformity using more extensible hydrogels that allow extensible filaments. We also measured the minimum possible diameter that can be obtained using each choice of hydrogel before breakage (**Figure 3e**). Analogous to previous findings, PTw80-1.5-1 achieved the smallest filament diameter reaching as low as 130µm. To further investigate the minimum filament diameter printed before disconnecting for PTw80-1.5-1, we measured the filament diameter at two additional higher value of *S* and obtained the average of 109 µm of filament thickness before disconnection (**Figure 3g and S11**). To summarize our findings, we compared the result of our work with current hydrogel based bioinks used in 3d extrusion bioprinting in the value of *S* and printed filament diameter (**Figure 3h**).^[9-10, 46-48]^ We show that the extensible HPMC-C_18_ hydrogel bioink can be extruded at a remarkably high value of *S* that is 5300-folds higher than the highest value of *S* reported for hydrogel-based bioinks tested with extrusion-based printing technology and achieve filament diameter 2-folds lower than any reported bioink by extrusion method. The comparison on print velocity with current works is also included, which shows that the improved speed index *S* is possible through controlling the flow velocity of the material (**Figure S12**). Here, we prove that HPMC-C_18_ hydrogel with the addition of Tw80 can improve the printability and print accuracy at high speed due to high filament extensibility.

## 3. Conclusion

We developed a tunable hydrogel platform by mixing hydrophobically modified cellulose-based polymers with various biomaterials. Our rheological studies demonstrated the tunability in viscoelastic properties and stress relaxation behavior without changing the stiffness through the incorporation of chemical additives, allowing selective control of rheological properties. We observed that the extensibility of the hydrogel was notably enhanced with the addition of Tw80 by FiSER and measured the extensional strain to break of up to 2000% in hydrogels with the highest Tw80 concentration. In 3D printing experiments, our hydrogels exhibited improved high-speed printability up to velocity factor increased over 5000-fold, mitigating issues of filament breakage and loss of uniformity. Our findings suggest that optimizing hydrogel extensibility through compositional adjustments offers promising enhancements for high-speed 3D printing, paving the way for fabricating complex, functional tissues with improved fidelity and efficiency.

## 4. Experimental Section

### 4.1 Materials

Hydroxypropyl methylcellulose stearoxy ether (Sangelose^®^ 90L) was provided from Daido Chemical Co. (Osaka, Japan). Tween 20 (polyethylene glycol sorbitan monolaurate), Tween 40 (polyethylene glycol sorbitan monopalmitate), Tween60 (polyethylene glycol sorbitan monostearate), Tween 80 (polyethylene glycol sorbitan monooleate), Tween 65 (polyoxyethylenesorbitan tristearate), Tween 85 (polyoxyethylenesorbitan trioleate), Span 20 (sorbitan monolaurate), Span 40 (sorbitan monopalmitate), Span 60 (sorbitan mono stearate), and alpha cyclodextrin were purchased from Sigma Aldrich and used as received.

### 4.2 HPMC-C_18_ Hydrogel Preparation

Hydroxypropyl methylcellulose stearoxy ether was prepared as a stock by dissolving in phosphate-buffered saline (PBS) at 6 wt %. The stock was left overnight to ensure that it was fully dissolved. Tween surfactants were prepared as a stock at 2wt% in PBS (except Tween 80 which was prepared at 8wt% in stock) and left overnight before use. Span surfactants were prepared at 30% in ethanol and heated using a heat gun until fully dissolved. αCD stock was prepared at 0.2wt% in PBS and left overnight before use. Hydrogel samples were prepared by thoroughly mixing the stock of HPMC-C_18_ with Tween, Span or alpha cyclodextrin and PBS in 1.5 ml Eppendorf tube using a spatula at a desired concentration. For example, to make 600mg of PTw20-1.5-0.5, 150mg of 6wt% stock of HPMC-C_18_, 150mg of 2wt% Tween 20 stock, and 300mg of PBS was added to 1.5ml Eppendorf tube and mixed well using a spatula. The tube was spun down at 10,000g for 5 minutes and mixed repeatedly until gelation occurred. All hydrogel samples were left overnight at room temperature prior to testing.

### 4.3 Shear Rheology

Rheological characterization was performed on a TA Instruments Discovery HR-2 stress-controlled rheometer using a 20mm diameter serrated plate with 500μm gap at 25°C. Solvent trap was used in all experiments to prevent dehydration. Frequency sweep measurements were performed at a constant 1% strain over frequencies from 0.1 rad.s^-1^ to 100 rad.s^-1^. Steady shear stress sweeps were performed from low to high with steady-state sensing and apparent yield stress values were defined as the stress at which the viscosity decreases 10% from the maximum. Amplitude sweep measurements were performed at a frequency of 0.1 rad/s over strain amplitudes of 1% to 10000% strain. P values were calculated with a one-way ANOVA followed by post hoc Tukey multiple comparison test unless otherwise noted.

### 4.4 Relaxation time fitting

The frequency sweep data were fit to a continuous relaxation spectrum framework based on the modified BSW spectrum to determine relaxation times from frequency sweep data where a clear crossover between storage and loss moduli was not observed in the frequency range studied. The data for *G*′ and *G*′′ were fit simultaneously to the spectrum using

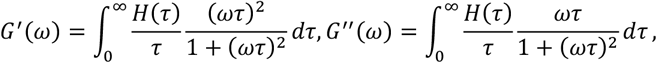

where *H*(*τ*) the modified BSW spectrum, suitable for modeling dynamic, glassy networks with incomplete/absent terminal regime relaxation^[49]^, is given by

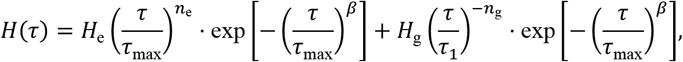

and the spectrum parameters were determined by least squares fitting of the combined residual using a differential evolution optimization algorithm on python. The relaxation timescale *τ*relax was determined from the fit spectrum using the following equation^[50, 51]^:

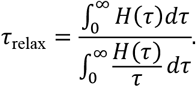

### 4.5 Extensional Strain-to-break Measurements

Extensional rheology was performed on a TA Instruments ARES-G2 rheometer in axial mode using an 8mm parallel plate geometry (R = 4mm) at a H = 4mm gap, resulting in an aspect ratio of H/R=1. Sample volume of 400 μL was loaded for each of the three replicated measurements. All experiments were performed at 25°C. Samples were loaded and tested immediately within seconds to minimize dehydration. Hencky (exponential) strain rates were applied at 0.06 s^-1^, 0.1 s^-1^ or 0.3s^-1^.

### 4.6 Cell culture, cell proliferation, and confocal imaging

NIH/3T3 fibroblasts (ATCC) were cultured in DMEM media with 10% FBS and 1% penicillin/streptomycin at 37° C and 5% CO2. For encapsulation, cells were trypsinzied with 0.25% trypsin/EDTA, rinsed, counted, and pelleted before being resuspended in PBS. They were then physically mixed with HPMC-C_18_/Tween mixtures to achieve the final material formulation. Materials with cell density of 5M/ml were cultured overnight at 37° C and 5% CO2. For cell proliferation assay, cell counting kit-8 (CCK-8, Sigma-Aldrich) was used to evaluate the proliferation of NIH/3T3 over a time course. Samples were cultured at 37°C and 5% CO2 in fresh culture media with 10% (v/v) CCK-8 solution in 96 well plate. Plate was read at 450 nm wavelength using Synergy H1 Microplate Reader (BioTek Instruments) to measure the optical density (OD). For imaging, live/dead stain (calcein/propidium iodide) was pipetted over the gel at 1M/ml and 1 hour was given for the dye to permeate the gel before imaging. Images were collected on a Leica confocal microscope.

### 4.7 3D Printing of Hydrogel Filaments

HPMC-C_18_ hydrogel was loaded in a BD 1ml syringe on a 27G needle for printing. Syringe-loaded hydrogel bioink was loaded onto the mechanically controlled extrusion-based 3D printer. All samples were printed at a constant speed of 350mm/min at varied flow rates. Pictures of printed filaments were taken immediately after the printing to prevent dehydration. Filament thickness measurements were done using ImageJ.

### 4.8 Tween 80 release assay

Glass capillary tubes were capped with epoxy at one end for the release assay of Tween 80 from the PTw80 hydrogels. PTw80 hydrogels (100μL) were injected into the bottom each tube and covered with 400μL of 1x PBS. Tubes were parafilmed to prevent dehydration and incubated at room temperature between timepoints. Samples were taken at each timepoint by removing 400μL of PBS without disturbing the gel surface and replaced with fresh PBS (400μl). Concentration of Tween 80 in each sample was measured using the flourescence enhancement property of probe 4′,5′-Dibromo-2′,7′-dinitrofluorescein disodium salt with the presence of Tw80.^[52]^

### 4.9 Statistical Analysis

For both shear rheology and extensional rheology measurements, sample size of 3 were tested. None of the samples was considered outliers in all experiments. For all experiments unless otherwise noted, comparison between multiple groups were conducted with a one-way ANOVA followed by post hoc Tukey multiple comparison test. Results were accepted as significant if p < 0.05. Values presented were means and standard deviations. For CCK-8 analysis and 3D printing success percentage, two-way ANOVA test is performed with Dunnett’s multiple comparison test with a single pooled variance. Values presented were means and standard deviations. Statistical testing was performed using Prism 10.

## Acknowledgements

This work was supported in part by the Bill & Melinda Gates Foundation (INV027411). Part of this work was performed at the Stanford Nano Shared Facilities (SNSF), supported by the National Science Foundation under award ECCS-2026822. We thank Mark Skylar-Scott group in Bioengineering department of Stanford University for providing 3D bioprinter used in this paper. O. M. S. is thankful for a National Science Foundation Graduate Research Fellowship and Hancock Fellowship of the Stanford Graduate Fellowship in Science and Engineering. C. K. J. is thankful for a National Science Foundation Graduate Research Fellowship.

## Author Contributions

Y.E.S., O.S., and E.A.A. conceived of the idea. Y.E.S., C.K.J and N.E. performed experiments and analyzed the data. Y.E.S., O.S. and S.S. analyzed the data.

## Materials and Data Availability

All data needed to evaluate the conclusions in the paper are present in the paper and/or the Supplementary Materials. Further information and requests for resources or raw data should be directed to and will be fulfilled by the lead contact, Eric Appel.

Received: ((will be filled in by the editorial staff))

Revised: ((will be filled in by the editorial staff))

Published online: ((will be filled in by the editorial staff))

