## Supplemental Information for "Highly Extensible Physically Crosslinked Hydrogels for High-Speed 3D Bioprinting"

**Supplementary Figures**

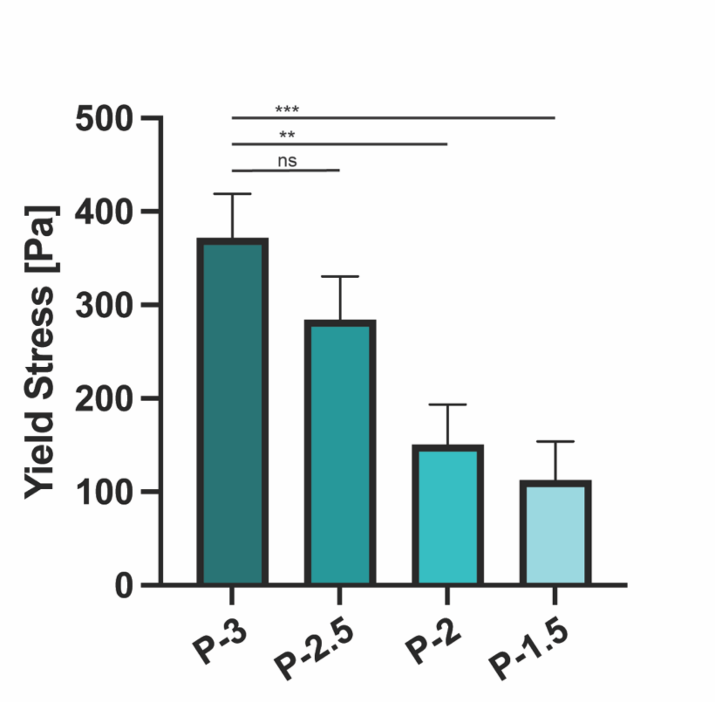


**Figure S1.** Yield stress of HPMC-$C_{18}$ hydrogel formulations. One-way ANOVA was used for multiple comparisons. Significance levels are indicated as $**p < 0.01$, and $***p < 0.001$.

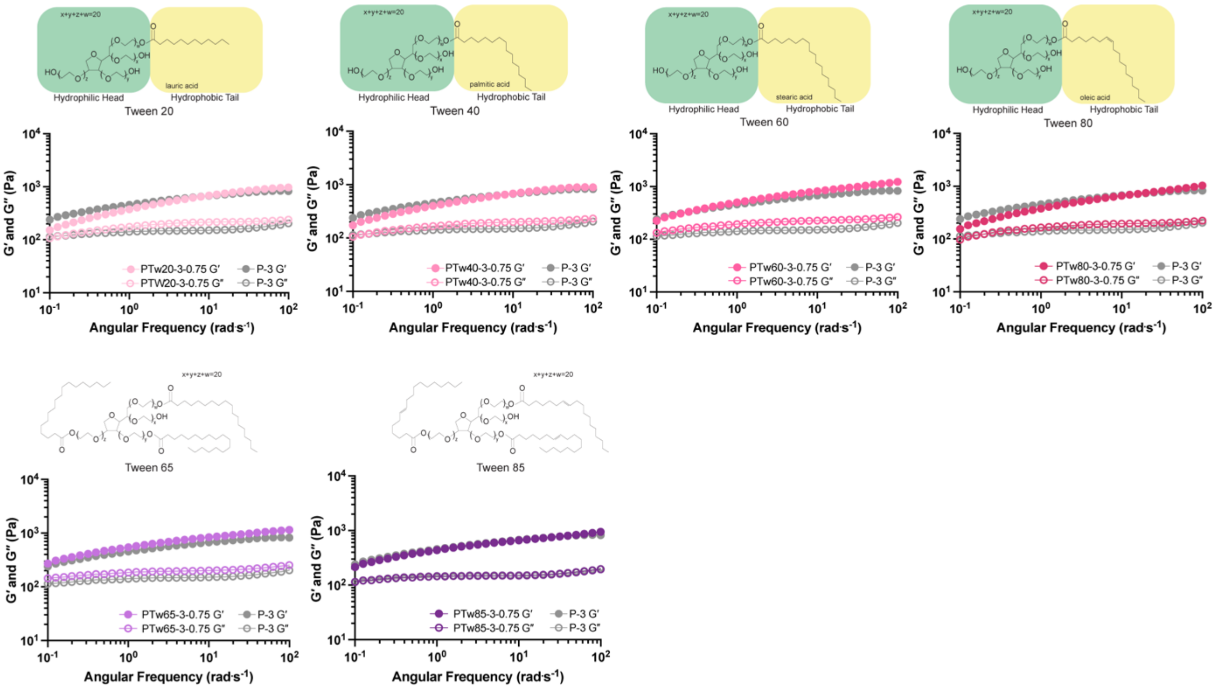


**Figure S2.** Frequency-dependent shear rheology of various HPMC-$C_{18}$ hydrogel formulations comprising different tween surfactants.

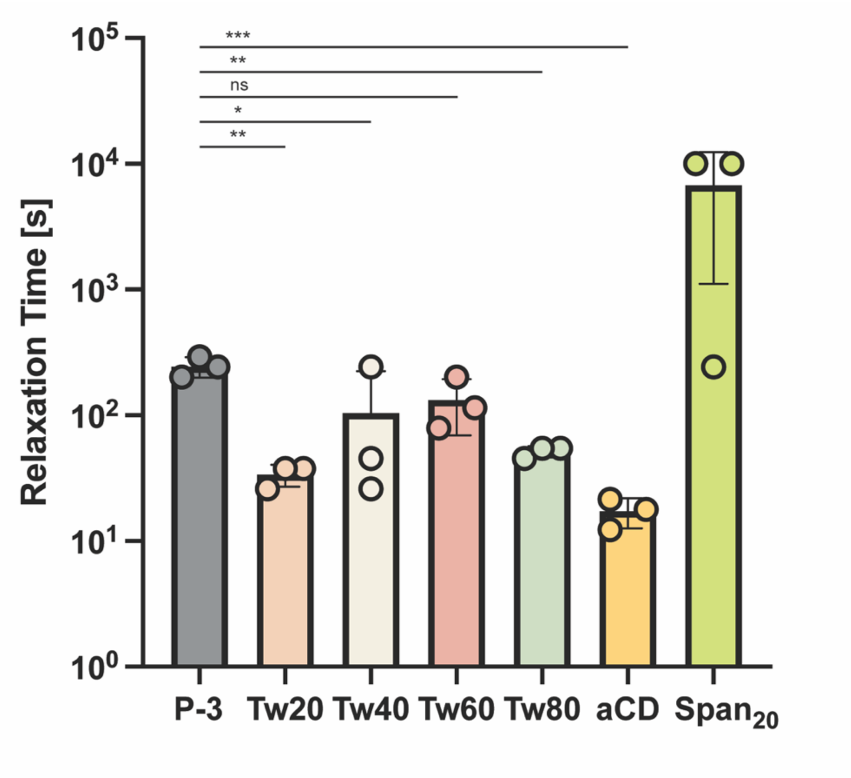


**Figure S3.** Relaxation time for various HPMC-$C_{18}$ hydrogel formulations comprising different tween surfactants. One-way ANOVA was used for multiple comparisons. Significance levels are indicated as $*p < 0.05$, $**p < 0.01$, and $***p < 0.001$.

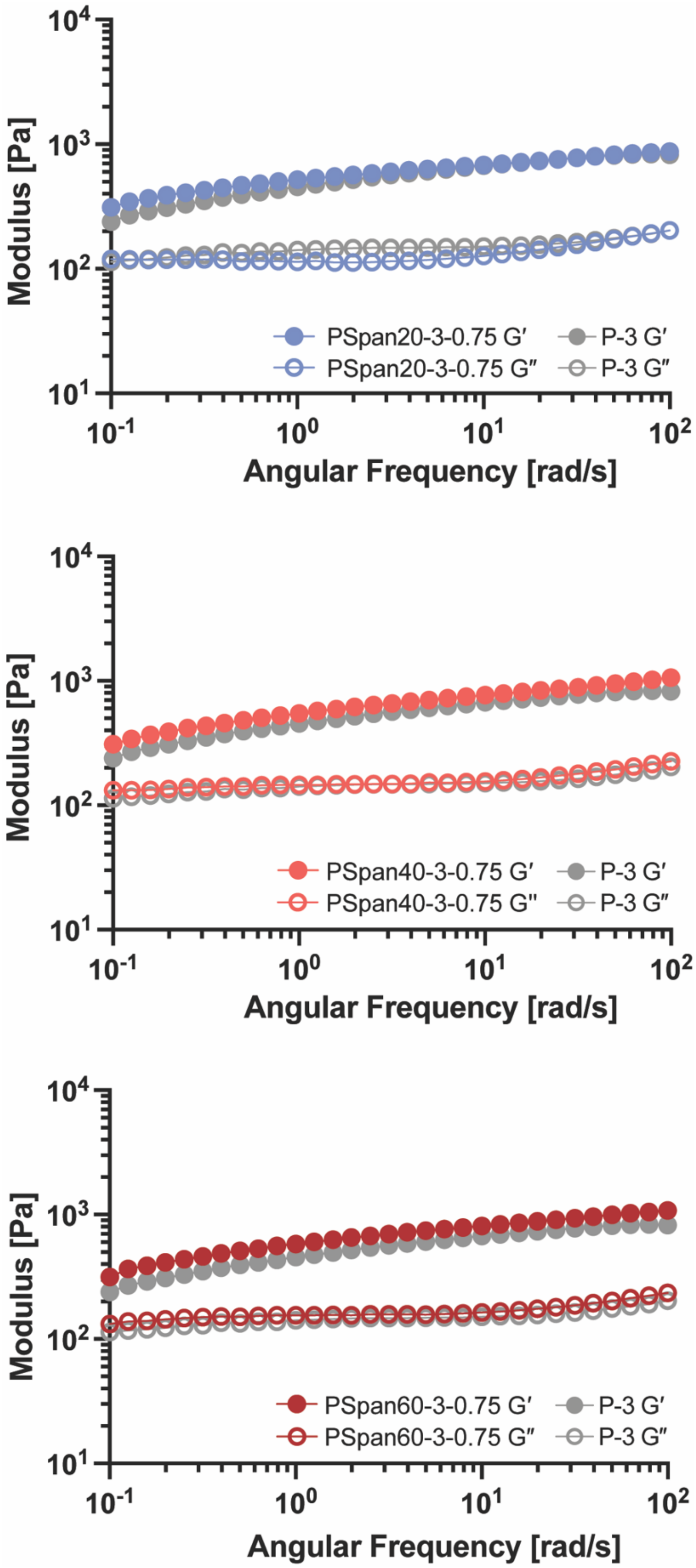


**Figure S4.** Frequency-dependent shear rheology of various HPMC-$C_{18}$ hydrogel formulations comprising different Span surfactants.

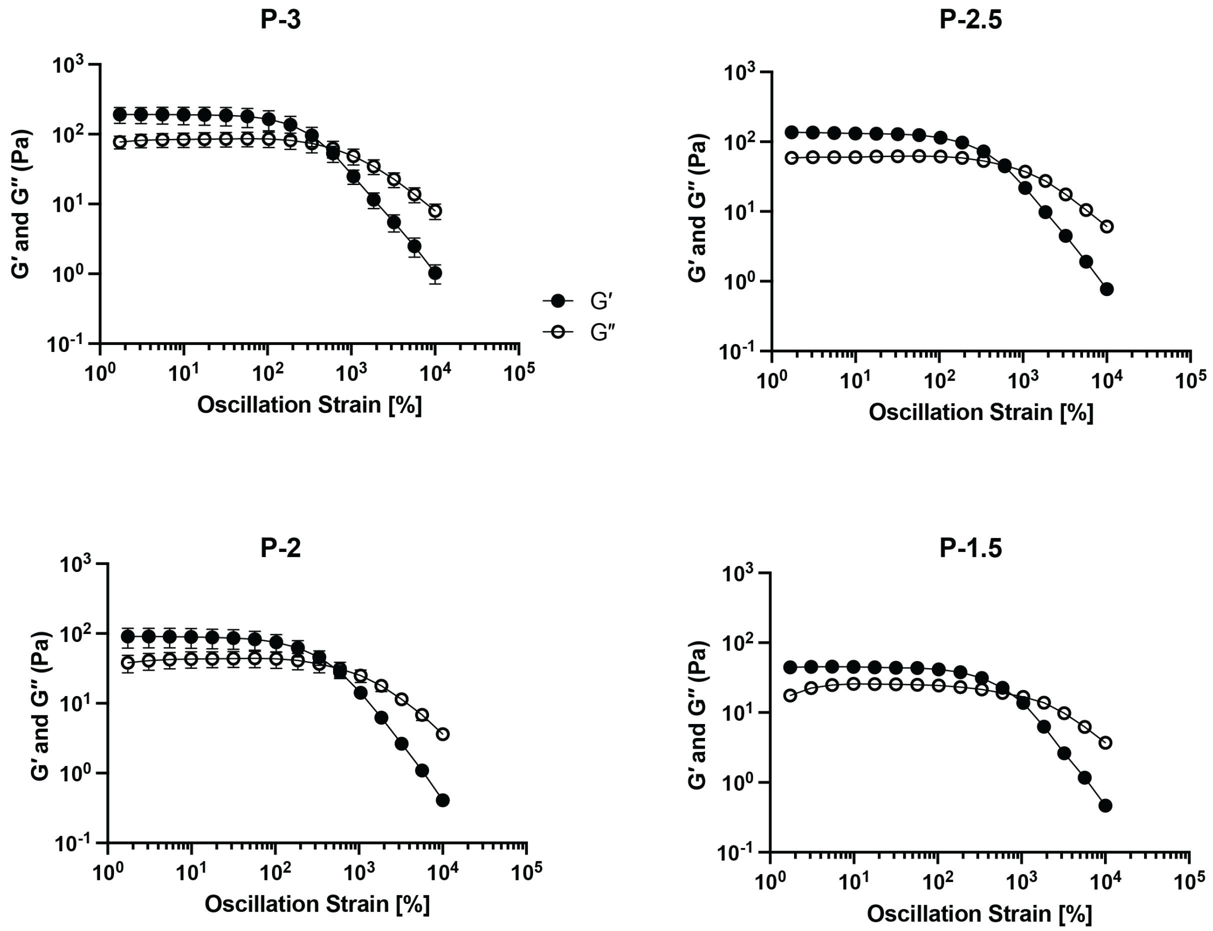


**Figure S5.** Amplitude-dependent shear rheology of various HPMC-$C_{18}$ hydrogel formulations.

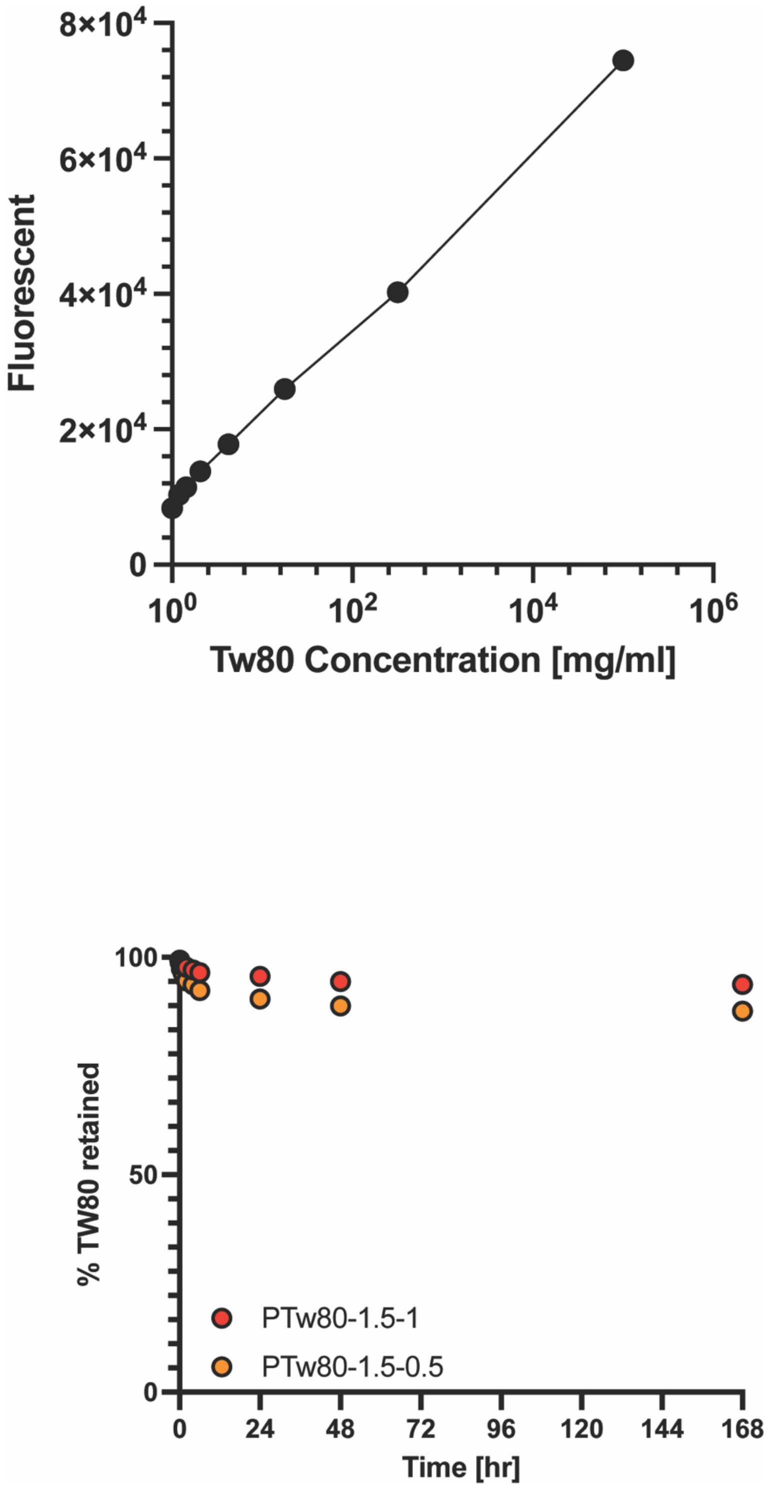


**Figure S6.** Retention of Tw80 in PTw80 hydrogels observed with *in vitro* release assays.

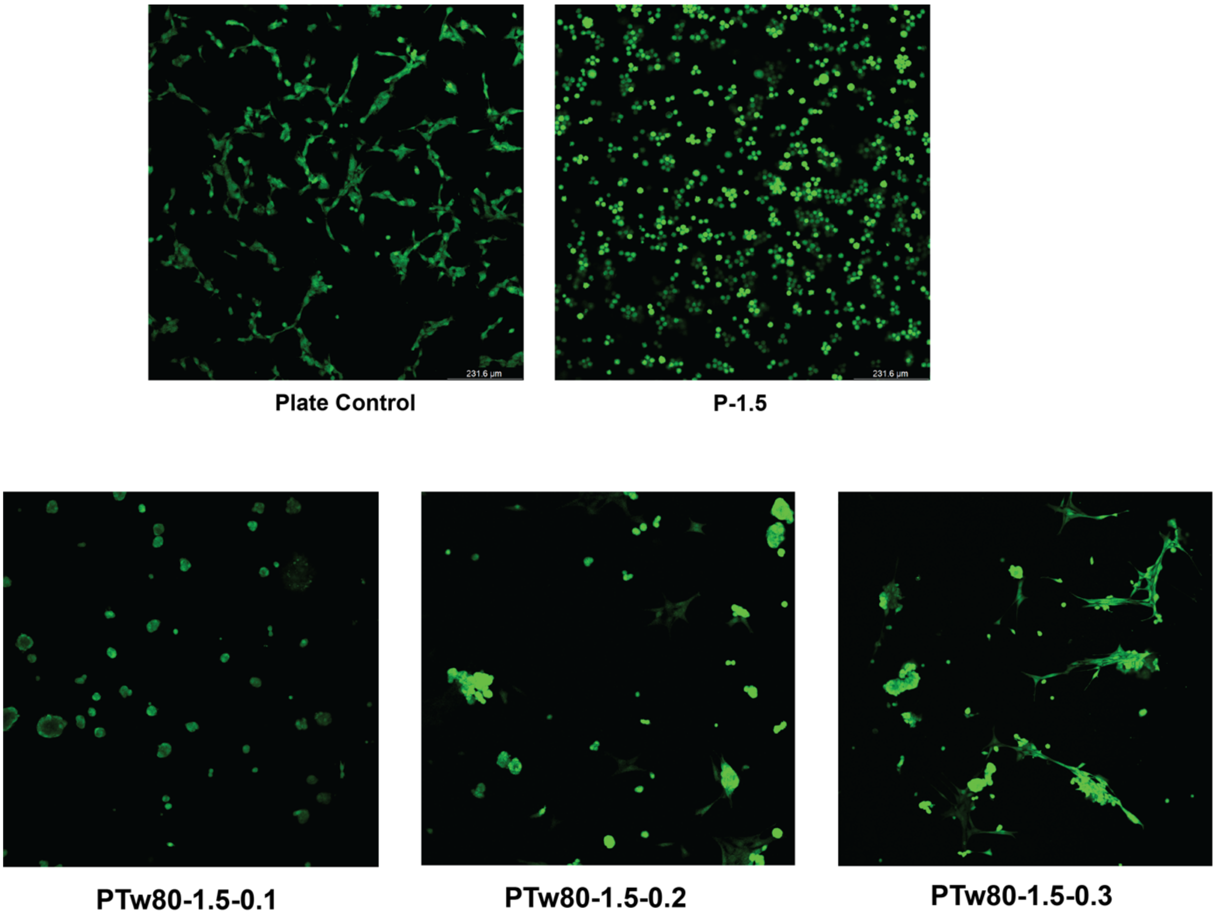


**Figure S7.** Confocal images of cells entrapped in PTw80 hydrogels.

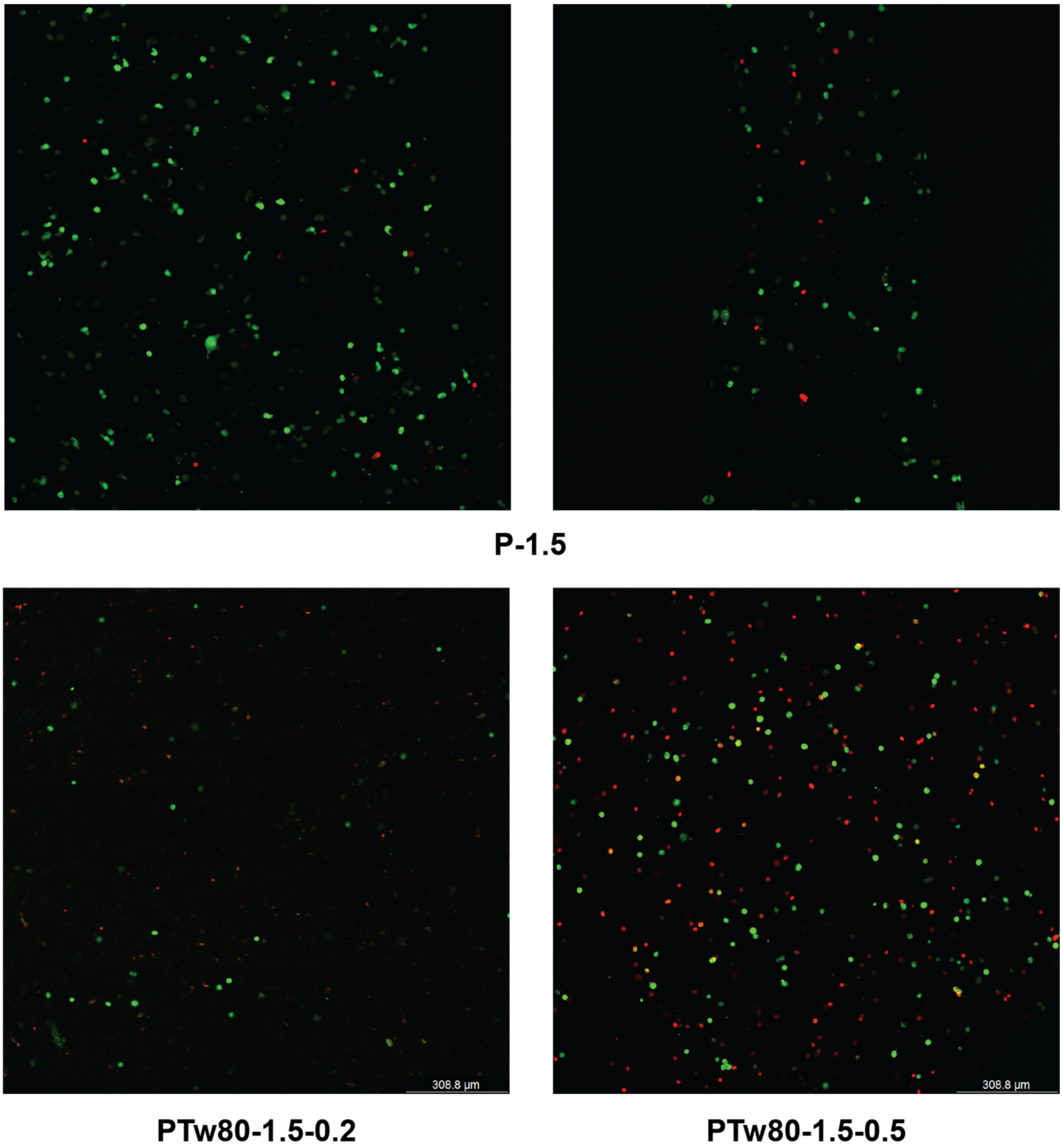


**Figure S8.** Confocal images of cells entrapped in PTw80 hydrogels comprising different amounts of Tw80 post printing.

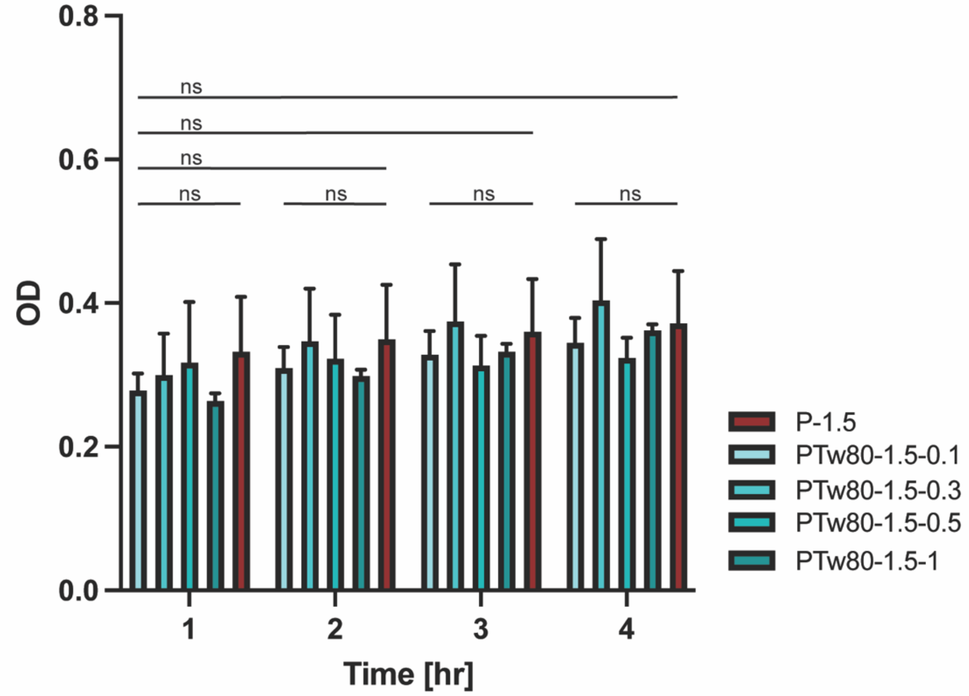


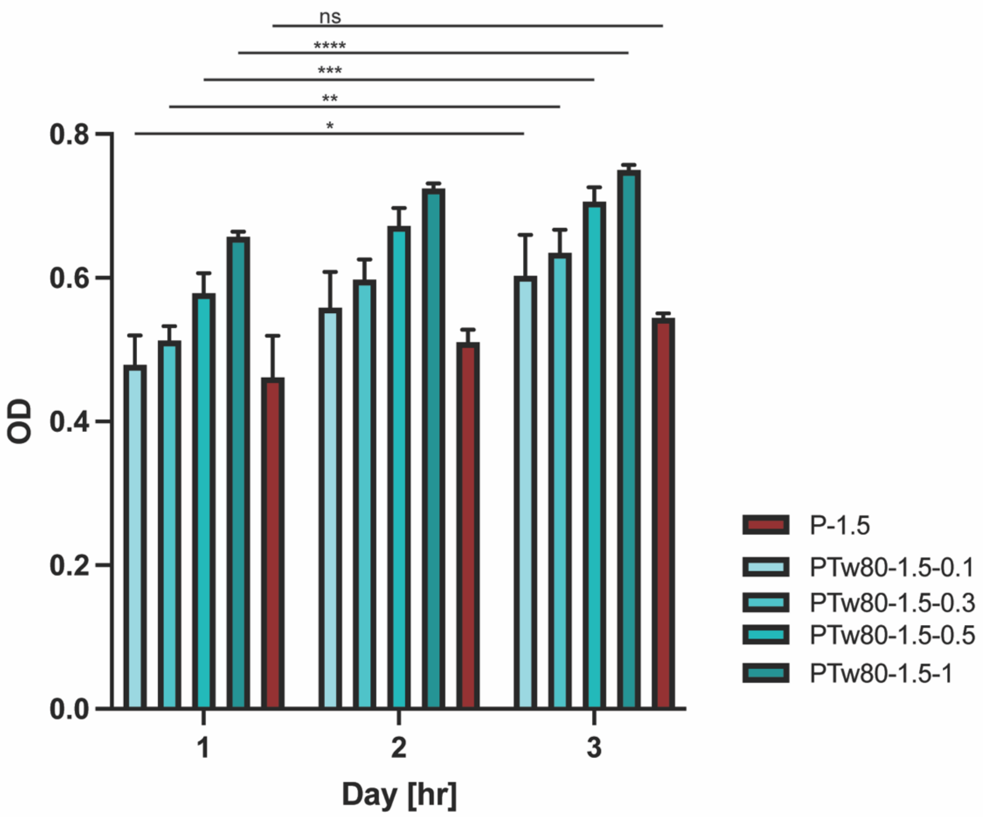


**Figure S9.** Cell counting kit-8 assay of PTw80 hydrogels. Two-way ANOVA was used for multiple comparisons. Significance levels are indicated as $*p < 0.05$, $**p < 0.01$, $***p < 0.001$, and $****p < 0.0001$.

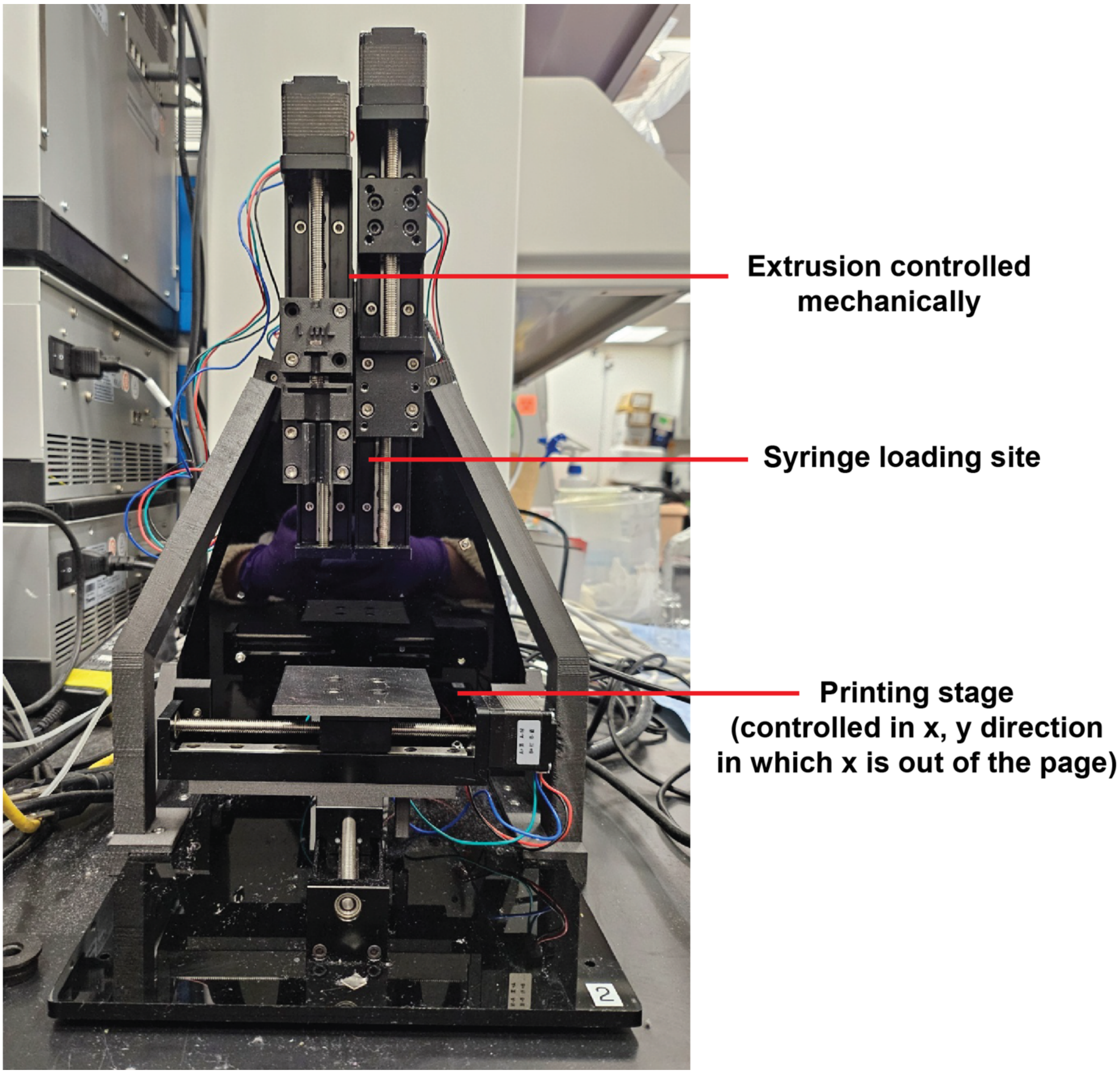


**Figure S10.** Extrusion-based 3D printer image.

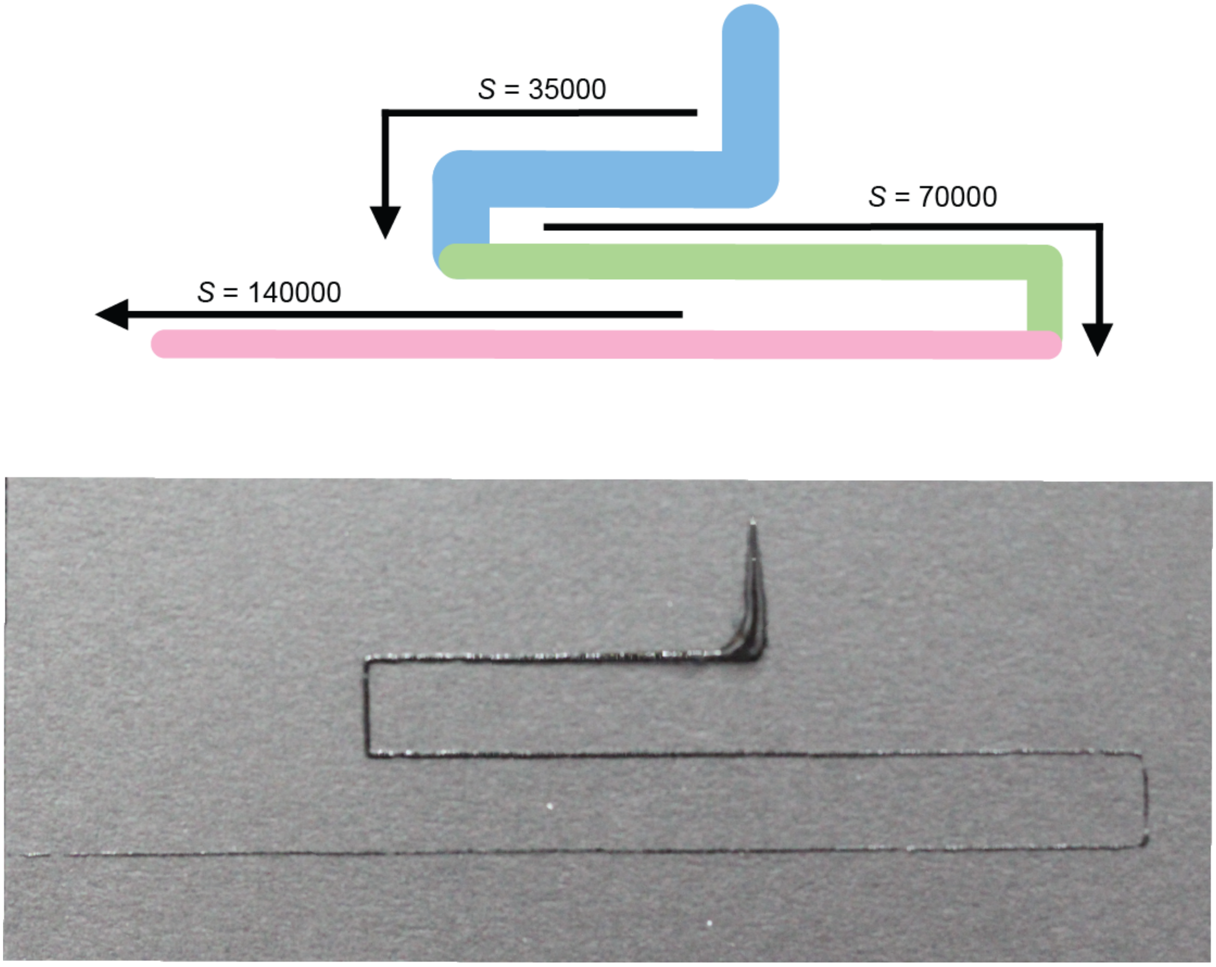


**Figure S11.** Schematic of 3d printing experiment for PTw80-1.5-1 and representative printed image.

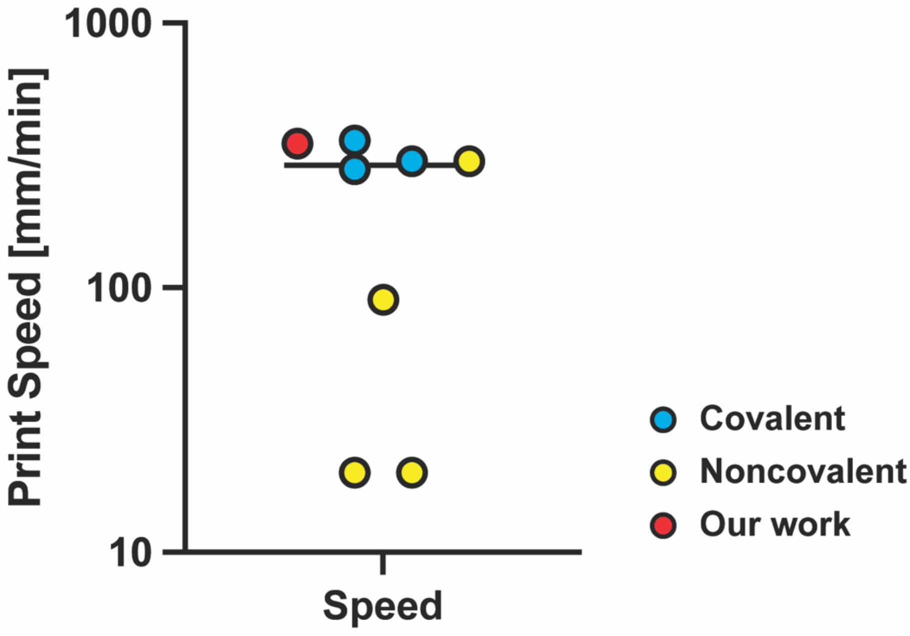


**Figure S12.** Absolute print speed (mm/min) comparison with hydrogel-bioinks in extrusion-based printing system reported in current literatures

**Supplementary Videos**

**Video S1.** Example filament stretching extensional rheometry (FiSER) of PTw80-1.5-1 at $1800 s^{-1}$.
